# Two antagonistic microtubule targeting drugs act synergistically to kill cancer cells

**DOI:** 10.1101/2020.02.06.936849

**Authors:** Lauralie Peronne, Eric Denarier, Ankit Rai, Renaud Prudent, Audrey Vernet, Peggy Suzanne, Sacnicté Ramirez-Rios, Sophie Michallet, Mélanie Guidetti, Julien Vollaire, Daniel Lucena-Agell, Anne-Sophie Ribba, Véronique Josserand, Jean-Luc Coll, Patrick Dallemagne, J. Fernando Díaz, María Ángela Oliva, Karin Sadoul, Anna Akhmanova, Annie Andrieux, Laurence Lafanechère

**Affiliations:** Institute for Advanced Biosciences, Team Regulation and Pharmacology of the Cytoskeleton, INSERM U1209, CNRS UMR5309, Université Grenoble Alpes, Grenoble, France; Grenoble Institute of Neurosciences, INSERM U1216, Université Grenoble Alpes, CEA, Grenoble, France; Cell Biology, Neurobiology and Biophysics, Department of Biology, Faculty of Science, Utrecht University, Padualaan 8, 3584 CH Utrecht, the Netherlands; NORMANDIE UNIV, UNICAEN, CERMN, Caen, France; Institute for Advanced Biosciences, Team Cancer targets and experimental therapeutics, INSERM U1209, CNRS UMR5309, Université Grenoble Alpes, Grenoble, France; Structural and Chemical Biology Department. Centro de Investigaciones Biológicas, CSIC, Ramiro de Maeztu 9, 28040 Madrid, Spain

**Keywords:** Cancer therapy, microtubules, drug synergy

## Abstract

Paclitaxel is a microtubule stabilizing agent and a successful drug for cancer chemotherapy inducing, however, adverse effects. To reduce the effective dose of paclitaxel, we searched for drugs which could potentiate its therapeutic effect. We have screened a chemical library and selected Carba1, a carbazolone, which exerts synergistic cytotoxic effects on tumor cells grown *in vitro*, when co-administrated with a low dose of paclitaxel. Carba1 targets the colchicine binding-site of tubulin and is a microtubule-destabilizing agent. The Carba1-induced modulation of microtubule dynamics increases the accumulation of fluorescent paclitaxel inside microtubules, providing a mechanistic explanation of the observed synergy between Carba1 and paclitaxel. The synergistic effect of Carba1 with paclitaxel on tumor cell viability was also observed *in vivo* in xenografted mice. Thus, a new mechanism favoring paclitaxel accumulation in microtubules can be transposed to *in vivo* mouse cancer treatments, paving the way for new therapeutic strategies combining low doses of microtubule targeting agents with opposite mechanisms of action.

## Introduction

Microtubules (MTs), dynamic polymeric filaments composed of α-tubulin and β-tubulin heterodimers, are key components of the cytoskeleton of eukaryotic cells. Their crucial roles in cell division and physiology mainly rely on their ability to rapidly polymerize or depolymerize. Targeted perturbation of this finely tuned process constitutes a major therapeutic strategy. Drugs interfering with MTs are major constituents of chemotherapies for the treatment of carcinomas. A number of compounds bind to the tubulin-MT system. They can be roughly classified into MT-stabilizers such as taxanes or epothilones, and MT-destabilizers such as vinca alkaloids, combretastatin and colchicine [1]. It has been demonstrated that binding of vinca alkaloids or colchicine prevents the curved-to-straight conformational change of tubulin at the tip of the growing MT, necessary for proper incorporation of new tubulin dimers into the MT lattice (see reviews [1,2]).

Paclitaxel (PTX) binds to the taxane-site of β-tubulin and stabilizes the MT lattice by strengthening lateral and/or longitudinal tubulin contacts within the MT [1]. At stoichiometric concentrations, it promotes MT assembly. At low and clinically relevant concentrations, PTX primarily suppresses MT dynamics without significantly affecting the MT-polymer mass [3,4]. PTX is one of the most successful chemotherapeutic drugs in history. It is currently used to treat patients with a variety of cancers including lung, breast and ovarian cancers [5].

Several mechanisms have been proposed to explain the anti-tumor activity of PTX. It can induce a mitosis dependent cell death, either by producing a mitotic arrest [6], when applied at high concentrations, or by promoting chromosome mis-segregation at low concentrations [7]. Alternatively, PTX can act on interphase cells and drive autonomous cell death by perturbation of intracellular trafficking [8]. It has also been recently proposed that post-mitotic formation of micronuclei induced by PTX can promote inflammation and subsequent tumor regression *via* vascular disruption and immune activation [9].

While PTX is a successful anti-cancer drug, its low solubility, its toxicity, and the fact that cells become resistant to this drug, impose serious limits to its use. Cell resistance to PTX is due to the high expression of P-glycoprotein or multidrug resistance-associated proteins, as well as to the overexpression of some β**-**tubulin isoforms or mutations in β-tubulin that affect the MT polymer mass and/or drug binding [10]. Another major drawback of PTX in clinical applications is the development of peripheral neuropathies, primarily involving the sensory nervous system. Although the molecular bases of these neuropathies are not completely understood, an inhibition of MT-based axonal transport appears to be a possible mechanism [11]. It has been recently shown that anterograde kinesin based-axonal transport is specifically affected by PTX, whereas MT destabilizing drugs that bind preferentially to the ends of MTs have much less effect on axonal transport [12].

An alternative therapeutic solution would be the use of pharmaceutics which, when co-administrated with PTX, could potentiate its effect without significantly increasing its toxicity. Such agents could allow the use of lower doses of PTX in cancer therapy, may limit the occurrence of resistances and reduce MT-independent adverse effects.

To identify such agents, we have screened a collection of 8,000 original compounds using a cytotoxicity assay and selected a derivative of the carbazolone series (Carba1) able to sensitize cells to a low, non-toxic dose of PTX. We demonstrated that Carba1 exerts synergistic cytotoxic effects with PTX. In cells, Carba1 has no major effect on the total MT mass in interphase cells and shows moderate cytotoxicity. We found that Carba1 targets the colchicine binding-site of tubulin and inhibits *in vitro* tubulin polymerization. Carba1-induced modulation of MT dynamics increases the accumulation of fluorescent PTX (Fchitax-3) inside MTs, providing a structural explanation of the observed synergy between Carba1 and PTX in cells.

Carba1 has no major anti-tumor effect when administrated alone in animals and no detectable toxicity. The administration of a combination of Carba1 and a low, ineffective, dose of PTX showed, however, a significant effect on tumor growth, indicating that Carba1 and PTX act synergistically *in vivo*. Our results pave the way for new therapeutic strategies, based on the combination of low doses of MT targeting agents with opposite mechanisms of action. These combinations may have reduced toxicity compared to high therapeutic PTX doses.

## Results

### A pairwise chemical genetic screen identifies a carbazolone derivative, Carba1, that sensitizes cells to paclitaxel

We designed a screen to select compounds that sensitize cells to paclitaxel (PTX). In a first step, we determined a minimal dose of PTX that is not toxic for cells. We found that 1 nM of PTX showed no toxicity when applied on HeLa cells for 48 hours **(Supplementary Fig 1)**. Furthermore, such a dose has no detectable impact on MT dynamics as assessed by EB3 tracking after time lapse fluorescence microscopy using GFP-EB3-transfected HeLa cells and subsequent calculation of dynamic instability parameters (**Supplementary Table 1**).

We then screened a library of 8,000 compounds at a concentration of 5 μM (**Fig 1A and Supplementary Table 2**) and compared their cytotoxicity on HeLa cells when administrated alone or in combination with 1 nM PTX. We selected 76 compounds that show no or moderate cytotoxicity when applied alone, and that were found cytotoxic when applied in combination with 1 nM PTX. We decided to focus our study on the 6-chloro-1,4-dimethyl-3-pyrrol-1-yl-9H-carbazole (Carba1, **Fig 1B**) because it did not display reactive chemical groups that could interact not specifically with protein targets and because it showed a synergistic activity with PTX (**Fig 1C**). Indeed, the comparison of HeLa cell apoptosis induced by Carba1 (12 μM), PTX (1 nM) to the apoptosis induced by the combination of Carba1 and PTX (12 μM/1 nM) confirmed the synergistic activity (**Fig 1D**).

**Figure 1:**
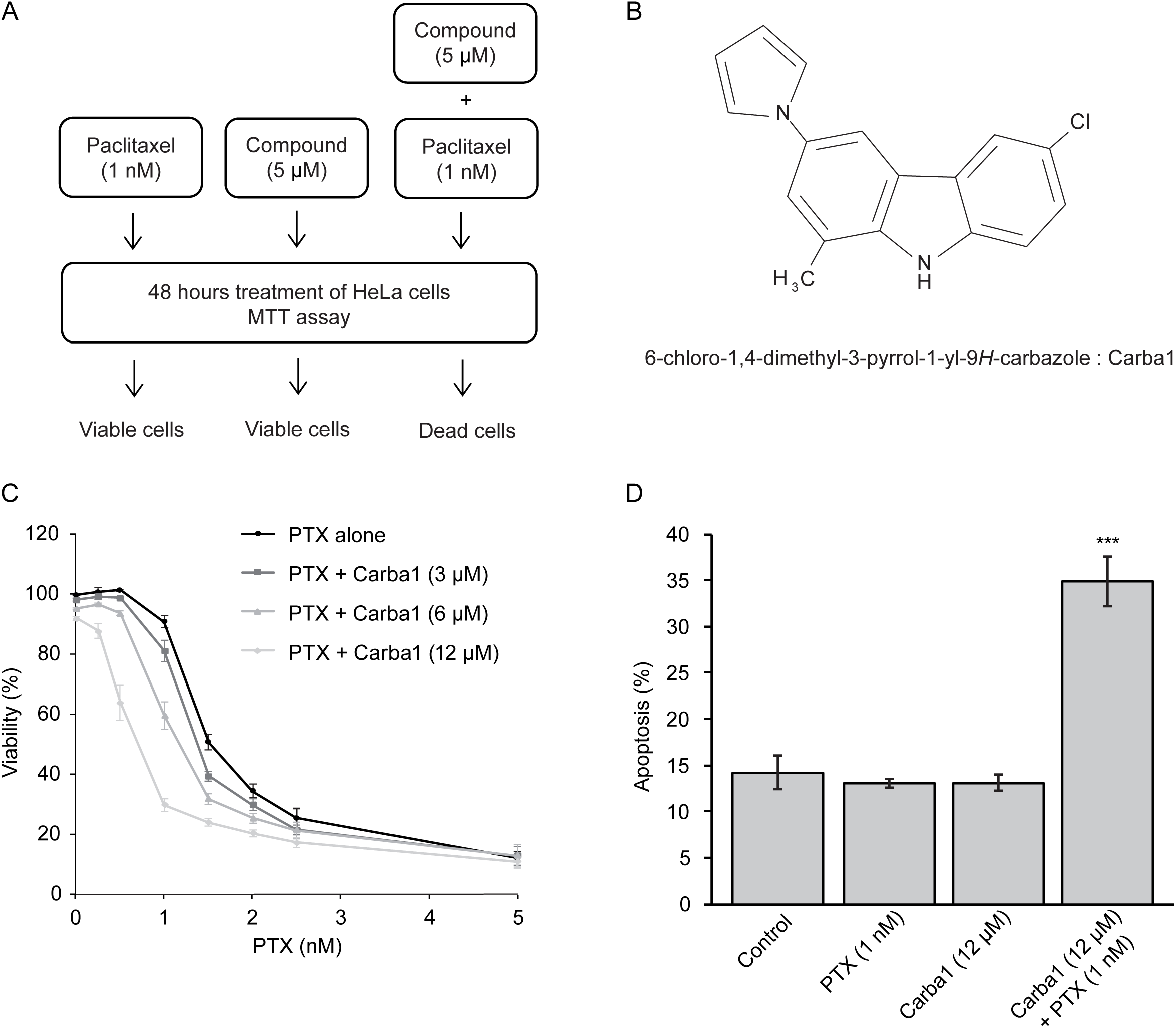
Selection of a compound that sensitizes cells to PTX. A Schematic illustration of the concept used to screen a chemical library for compounds that sensitize cells to PTX. Treatment of cells with compounds of the library alone (5 μM) or PTX (1 nM) alone has no effect on cell viability, compounds (5 μM) that have no effect when applied alone but induced cell death when applied in combination with PTX (1 nM) were selected. B Chemical structure of Carba1. C Effect of Carba1/PTX combinations on the viability of HeLa cells. Cells were incubated for 72 hours with the indicated combinations of Carba1/PTX. The percentage of viable cells was calculated following a Prestoblue assay. Data are presented as mean ± SEM of 3 independent experiments. D Effect of Carba1 (12 μM), PTX (1 nM) and the combination of Carba1 and PTX (12 μM/1 nM) on HeLa cell apoptosis. HeLa cells, treated with the indicated concentrations of drugs, were stained with propidium iodide and annexin V and analyzed by flow cytometry. Results are expressed as mean ± SEM of 3 separate experiments. The significance was determined by a Student’s t-test (***p<0.001, compared to the control).

### Carba1 has a moderate cytotoxicity when applied at high concentrations

As our final aim was to test the therapeutic efficacy of Carba1 in combination with PTX, it was important to investigate its cellular effects and to check that this compound is not or moderately toxic by itself. We first analyzed the cytotoxicity of Carba1 on HeLa cells, using the sensitive PrestoBlue assay. As shown in **Fig 2A**, Carba1 has a moderate cytotoxicity with a calculated GI50 of 21.8 μM after a 72-hour treatment.

**Figure 2:**
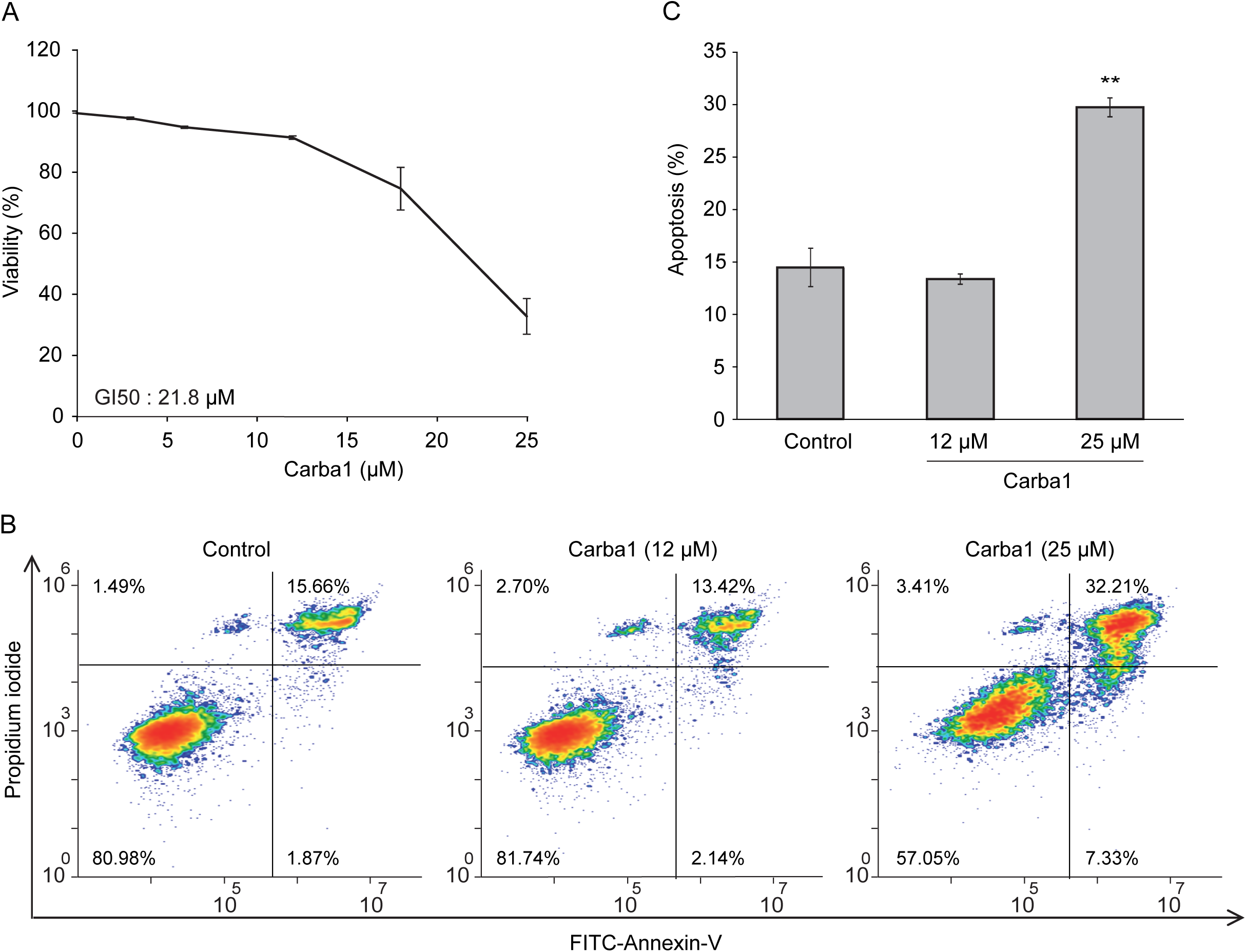
Analysis of Carba1 toxicity. A Effect of Carba1 on HeLa cell viability. Cells were incubated for 72 hours with increasing concentrations of Carba1. The percentage of viable cells was calculated following the Prestoblue assay. The results are expressed as mean ± SEM of three separate experiments. B Effect of Carba1 on HeLa cells apoptosis. HeLa cells, treated with the indicated concentrations of Carba1 for 72 hours, were stained with propidium iodide and FITC-annexin V and analyzed by flow cytometry. Apoptotic cells are observed in the upper right part of the graphs. C Results for apoptotic cell death (as shown in figure 2B) are expressed as mean ± SEM of 3 separate experiments. The significance was determined by a Student’s t-test (**p<0.01, compared to the control).

Since the Prestoblue assay is a metabolic test that indirectly measures cell viability, we directly detected cells in apoptosis using Annexin V staining, and quantified them by flow cytometry. We compared the effect of two concentrations of Carba1: a concentration (12 μM) that has no detectable effect on cell viability and a cytotoxic concentration (25 μM). No apoptosis was detected when Carba1 was applied for 72 hours at a concentration of 12 μM whereas at 25 μM, it induced apoptosis of 30% of the cells (**Fig 2B and 2C**).

These results indicate that Carba1 is only weakly toxic, even when applied at a high concentration. A toxicity analysis of a single 10 μM dose of Carba1 on a set of 60 human cancer cell lines (NCI-60 screen [20]) confirmed the low cytotoxic activity of Carba1 (**Supplementary Table 3**).

### Cell-cycle progression is blocked at mitosis by Carba1

A videomicroscopy analysis, using different doses of Carba1, showed that the compound impacted mitosis. As compared to DMSO, Carba1 (12 μM) induced a significant delay in the completion of metaphase and a slight increase of aberrant mitosis (**Fig 3A, B, Supplementary Movie 1 and Supplementary Table 4)**. When Carba1 was applied at a concentration of 25 μM, the majority of the cells stayed blocked in prometaphase (**Fig 3A, B** and **Supplementary Movie 1, right**). We followed and quantified the fate of the cells treated with 25 μM Carba1 in a 20-hour time lapse video (**Supplementary Movie 2**) and noted that 61% of the mitotic cells eventually died during mitosis, 29% were still dividing abnormally, whereas only 10% underwent apparently normal mitosis (**Supplementary Table 4**). We thus concluded that a cytotoxic dose of Carba1 induced a very long duration of mitotic arrest, followed by mitotic catastrophe.

**Figure 3:**
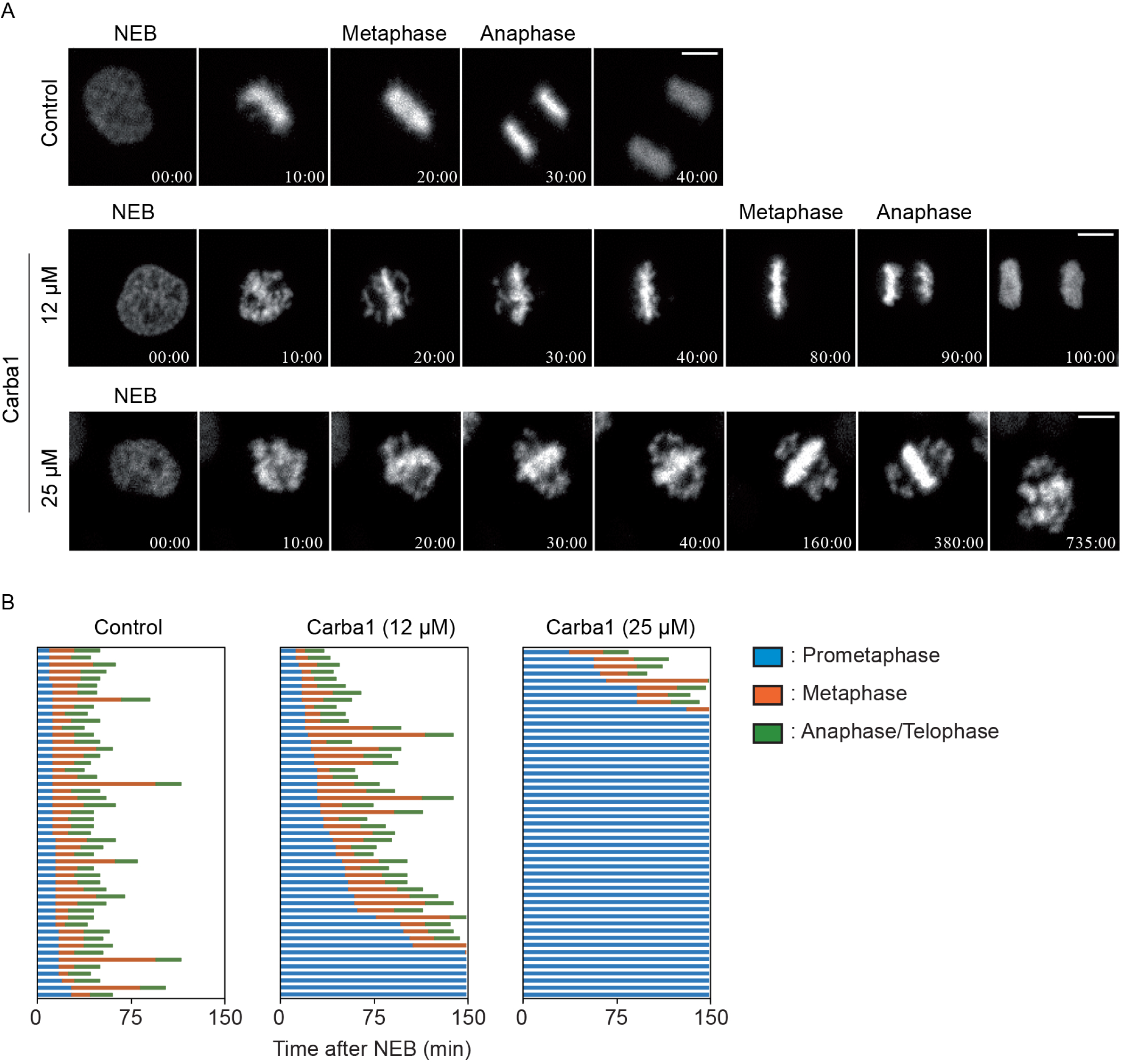
Carba1 induces a mitotic arrest. A Representative images, selected from supplementary movie 1 of HeLa Kyoto cells treated with DMSO (control) and the indicated concentrations of Carba1. Bar =10 μm. B Analysis of the duration of mitosis in HeLa Kyoto cells treated with DMSO (control) or with different doses of Carba1, as indicated. Duration of prometaphase (from nuclear envelope breakdown (NEBD) to chromosome alignment; blue), metaphase (from chromosome alignment to anaphase onset; orange) and anaphase/telophase (from anaphase onset to chromosome decondensation; green) were analyzed from supplementary movie 1. The data represent 50 cells for each treatment.

In accordance with the effect of Carba1 on mitosis, a flow cytometry analysis using propidium iodide staining indicated that a 15-hour exposure to 25 μM Carba1 induced a dose-dependent cell-cycle arrest at the G2/M phase (**Fig 4A**). Prolonged exposure (24 and 48 hours) led to a reduction of the number of cells blocked in the G2/M phase and to an increase of aneuploid cells, as assessed by the increased number of cells in sub G1 and of cells containing more than 4N DNA (**Fig 4A**).

**Figure 4:**
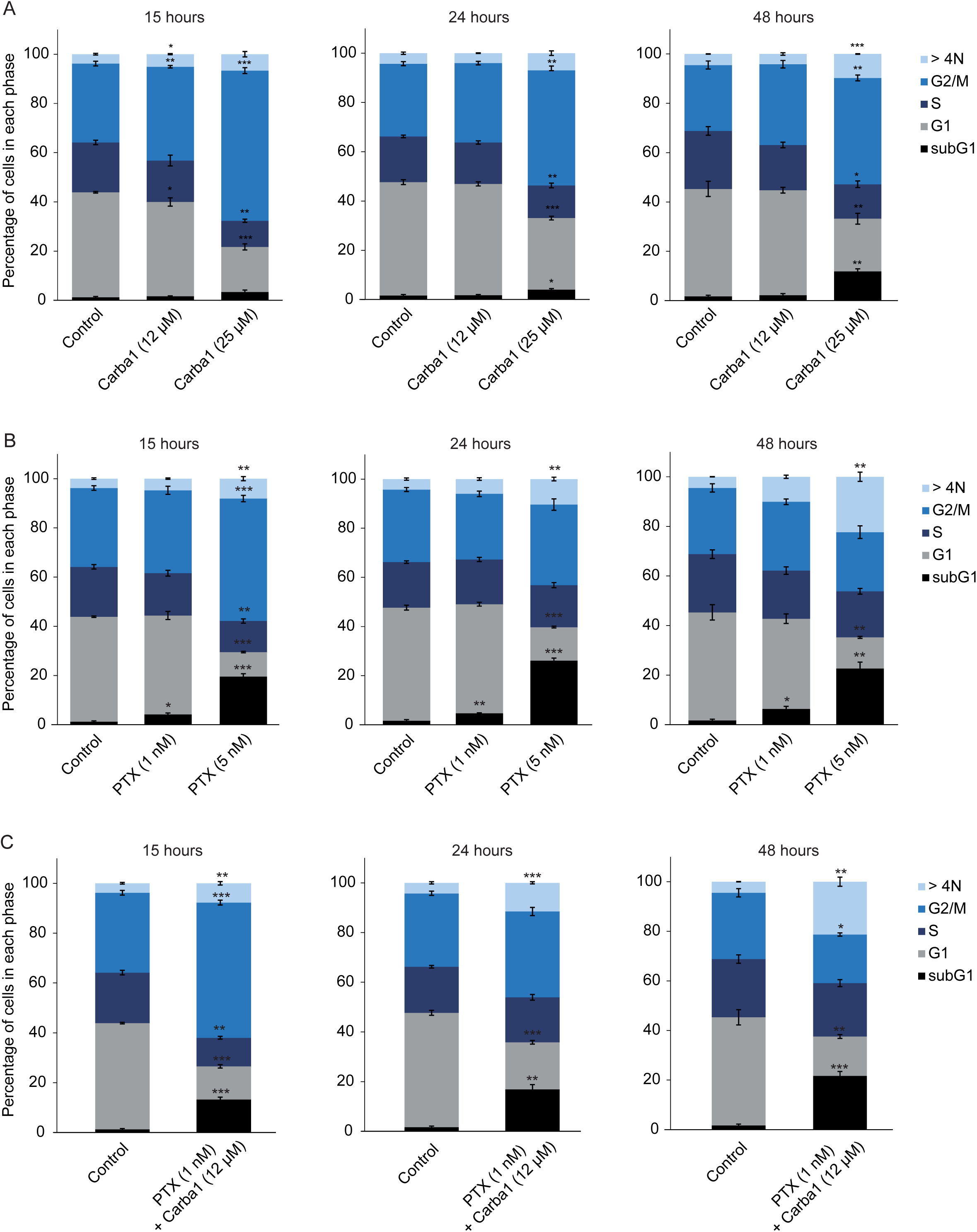
Comparative analysis of the effect of Carba1, PTX and of a Carba1/PTX combination on the cell cycle. A HeLa cells treated with the indicated concentrations of Carba1 for 15, 24 and 48 hours, were fixed with 70% ethanol, stained with propidium iodide and analyzed by flow cytometry. B HeLa cells treated with the indicated concentrations of PTX for 15, 24 and 48 hours, were fixed with 70% ethanol, stained with propidium iodide and analyzed by flow cytometry. C HeLa cells treated with the combination of Carba1 and PTX (12 μM/1 nM) for 15, 24 and 48 hours, were fixed with 70% ethanol, stained with propidium iodide and analyzed by flow cytometry. The results are expressed as mean ± SEM of 3 separate experiments. The significance was determined by a Student’s t-test (*p<0.05, **p<0.01, ***p<0.001, compared to the control).

### Carba1 increases PTX effects on cell cycle and mitosis

We similarly analyzed the effect of PTX, using time-lapse microscopy. PTX at a concentration of 1 nM induced a delay in chromosome congression during prometaphase and a moderate increase of aberrant mitosis (**Supplementary Movie 3, left** and **Supplementary Table 4).** When treated with a cytotoxic concentration of PTX (5 nM) 80% of HeLa cells underwent aberrant mitosis followed by a mitotic slippage, as shown by a 12-hour time-lapse video (**Supplementary Movie 3, middle** and **Supplementary Table 4**). We conducted a flow cytometry analysis to get further insight of the effect of 5 nM PTX on the fate of HeLa cells treated for longer times (15, 24 or 48 hours). After a 15-hour treatment, half of the cell population was blocked in the G2/M phase and nearly 20% of the cells were dead, as indicated by the increased proportion of cells in subG1. Then, the proportion of cells in G2/M gradually decreased, in parallel with an increased number of cells in subG1 and of plurinucleated cells (**Fig 4B**). Because the effects of such a cytotoxic concentration of PTX were different from those of a cytotoxic (25 μM) concentration of Carba1, we wondered which compound effect was predominant in the cytotoxicity of the combination of Carba1 and PTX (12 μM/1 nM). We thus compared the effects of this cytotoxic combination to the effects of Carba1 25 μM and PTX 5 nM administrated separately. As shown on **Fig 4C**, the combination of Carba1 and PTX (12 μM/1 nM) induced an arrest of the cell cycle almost superimposable to the arrest observed when cells are treated with PTX 5 nM. Moreover the videomicroscopy analysis of the cells treated with this combination showed that cell death occurred after mitotic slippage (**Supplementary Movie 3, right**). The similarity of the results obtained with the combination Carba1 and PTX (12 μM/1 nM) to those obtained with PTX 5 nM indicates that the overall effect of the combination results from an increase of the PTX effect induced by Carba1.

### Carba1 is a microtubule-destabilizing agent

In an attempt to understand the Carba1 mechanism of action, we first analyzed its effect on cellular MTs, using immunofluorescence. Carba1 treatment (12-25 μM) did not visibly perturb the MT network in interphase cells, as compared to DMSO (control; **Fig 5A**). In mitosis, chromosome congression defects were visible in several mitotic cells of the 12 μM treated cell population. The occurrence of such defects was increased at a higher dose (25 μM) of Carba1 (**Fig 5A**). Such defects in chromosome congression are similar to those observed on cells treated by some inhibitors of kinases involved in the mitotic process such as Aurora B or Plk1 kinases [21]. Moreover, compounds structurally related to Carba1 often target protein kinases [22,23]. We therefore tested the ability of Carba1 to inhibit the activity of a panel of 64 protein kinases including kinases known to be involved in the regulation of the cytoskeleton and/or the cell cycle. We found that, when *in vitro* assayed at a 10 μM concentration, Carba1 did not show any selective inhibitory activity on the kinases tested (**Supplementary Table 5**). It is therefore unlikely that Carba1 is a direct inhibitor of these kinases.

**Figure 5:**
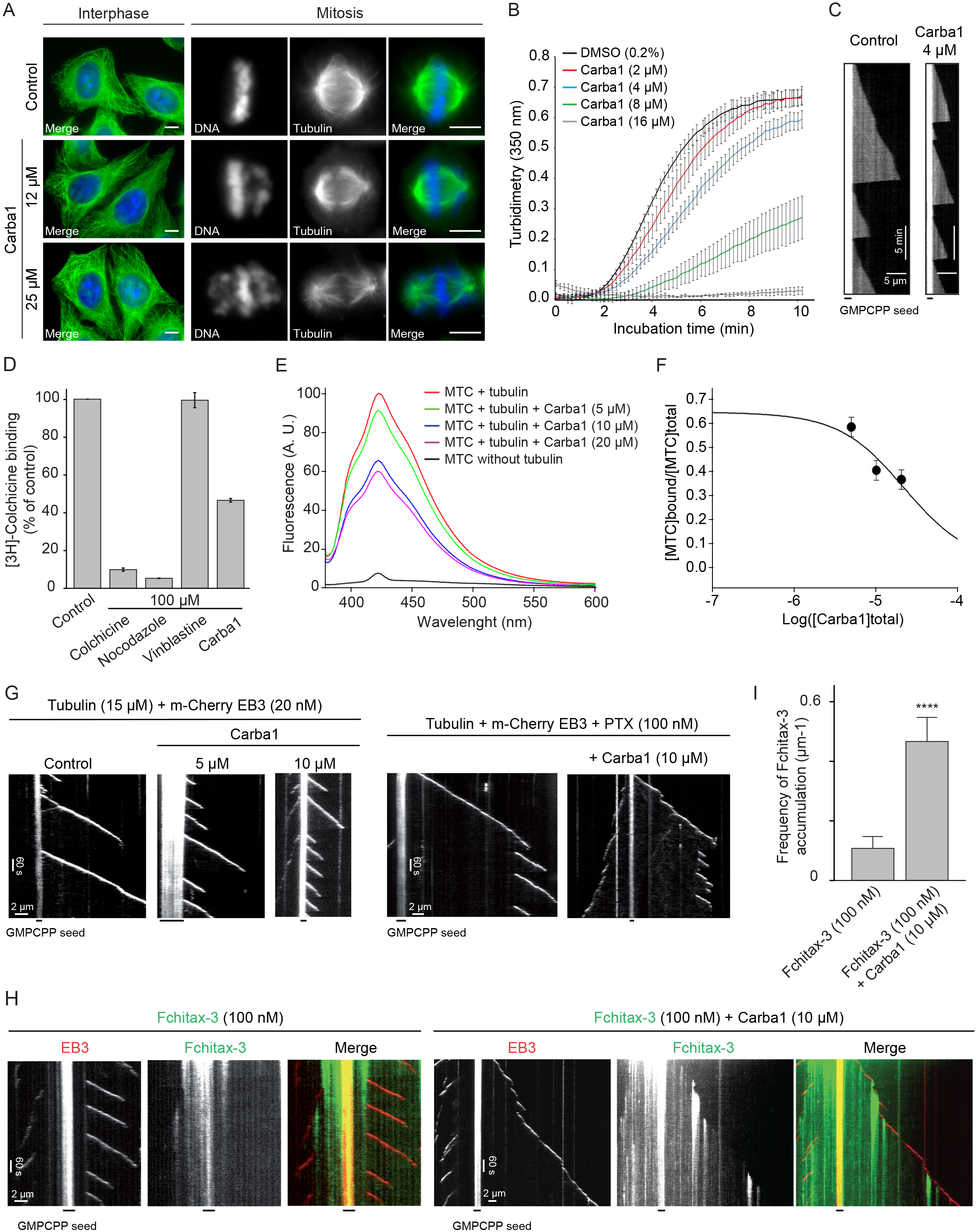
Effect of Carba1 and of the Carba1/PTX combination on MTs. A Immunofluorescence analysis of the Carba1 effect on interphase and mitotic MTs. MTs in interphase (left panels) or in mitosis (right panels) were stained using an anti-tubulin antibody, as described in the methods section. DNA was stained using Hoechst reagent. Bars = 10 μm. B Time course of tubulin polymerization at 37°C in the presence of vehicle (DMSO, black line) and Carba1 at different concentrations (colored lines) as indicated, measured by turbidimetry at 350 nm. Purified tubulin: 30 μM in BRB80 buffer with 1 mM GTP. Each turbidity value represents the mean ± SEM from 3 independent experiments. C 12% ATTO 488-labeled MTs (12 μM free tubulin dimers) were grown from MT seeds stabilized by GMPCPP in the presence of different concentrations of Carba1 on a cover glass and then detected by TIRF microscopy. Representative kymographs for control and 4 μM Carba1 conditions, illustrating MT plus end growth. D Effect of Carba1 on the binding of [^3^H]-colchicine to tubulin. Carba1 (100 μM) was used to compete with [^3^H]-colchicine (50 nM) as described in the methods section. Each value represents the mean ± SEM of 3 independent experiments. Colchicine and nocodazole were used as positive and vinblastine as negative control. E Displacement of MTC from the colchicine site. Fluorescence emission spectra of 10 μM MTC and 10 μM tubulin in 10 mM phosphate-0.1 mM GTP buffer pH 7.0, in the absence or presence of increasing concentrations of Carba1. F Displacement isotherm at 25°C of the fluorescent probe MTC (10 μM) bound to tubulin (10 μM) by Carba1 (black line and circles). The solid line is the best-fit value of the binding equilibrium constant of the competitors, assuming a one-to-one binding to the same site. G, H Kymographs illustrating MT plus end growth in the presence of 15 μM tubulin, 20 nM m-Cherry EB3 without (control) or with 5 and 10 μM Carba1, PTX (100 nM) without or in combination with 10 μM Carba1, Fchitax-3 (100 nM) without or with 10 μM Carba1. I Quantification of Fchitax-3 accumulation frequencies per MT unit length in the presence of 15 μM tubulin with 20 nM mCherry-EB3 without (n=11) or with 10 μM Carba1 (n=13). Each value represents the mean ± SD of 2 independent experiments.

The observed effects of Carba1 on the cellular MT network were reminiscent to those described for low doses of MT depolymerizing agents such as nocodazole or vinca alkaloids: a mitotic arrest with a similar aberrant chromosome organization, with no detectable effect on the total MT mass [24]. We thus wondered if Carba1 was, as nocodazole, able to directly impact MT assembly. The effect of a high dose (25 μM) of Carba1 on MT dynamic instability parameters was measured using time-lapse fluorescence microscopy on GFP-EB3 transfected cells (**Supplementary Table 1**). Carba1 reduced the MT growth rate as well as the MT growth length, as indicated by the increase of the distance-based catastrophe frequency, and increased time spent in pause, indicating that Carba1 suppresses MT dynamics.

We therefore tested the Carba1 effect on *in vitro* tubulin assembly. As shown in **Fig 5B**, Carba1 was able to inhibit polymerization of pure tubulin in a dose-dependent manner. Increasing Carba1 doses induced a decrease in the rate of polymerization, as well as a delay in nucleation and a reduction in the total quantity of assembled MTs, attested by the scaling down of the level of assembly at equilibrium (**Fig 5B**). The concentration of Carba1, which inhibits 50% of tubulin (30 μM) assembly under these experimental conditions, was 6.9 μM. To assay the effects of Carba1 on MT dynamics more accurately, we assembled fluorescently labeled, dynamic MTs *in vitro* and used total internal reflection fluorescence (TIRF) microscopy to track individual MTs (**Fig 5C**). We found that 4 μM Carba1 had no effect on either MT catastrophe (switch from growth to shrinkage) or rescue (switch from shrinkage to growth) frequencies, but induced a significant reduction in MT growth rates. Moreover, the time spent inactive before a new polymerization was largely increased for Carba1 treated MTs (**Supplementary Table 6**).

We then looked for the binding site of Carba1 on tubulin. Among the four binding sites described for MT depolymerizing agents, the most common binding site is the colchicine site [1]. We checked whether Carba1 can compete with [^3^H]-colchicine for its tubulin binding-site (**Fig 5D**). Carba1 selectively inhibited colchicine binding to tubulin, indicating that it binds to tubulin at or near the colchicine site. However, it did not completely prevent the binding of [^3^H]-colchicine, suggesting that its affinity for this site is lower than that of colchicine.

In order to measure the binding constant of the compounds, a competition assay with 2-methoxy-5-(2,3,4-trimethoxyphenyl)-2,4,6-cycloheptatrien-1-one (MTC), an analogue of colchicine lacking the B ring that rapidly reaches an equilibrium (Kb= 4.7×10^5^ M^−1^, 25°C [25]) in its binding reaction with tubulin, was designed. In the absence of tubulin the compound lacked fluorescence (**Fig 5E**) while in the presence of tubulin an emission maxima at 423 nm was observed upon excitation at 350 nm. As expected from its activity as an inhibitor of [^3^H]-colchicine binding to tubulin, Carba1 is able to displace MTC from the colchicine site, strongly supporting that Carba1 binds to the colchicine site of tubulin. The dissociation constant of Carba1, for the colchicine site is 3.03 ± 0.5 ×10^−6^ mol L-1 (**Fig 5F**). Altogether, these results show that Carba1 is a direct inhibitor of MT polymerization.

### Carba1 binding to tubulin enhances the tubulin binding capacity of PTX and its MT stabilizing activity

Carba1 is thus a compound that binds tubulin and impairs MT polymerization. We wondered how such a mechanism of action could explain the synergy between Carba1 and PTX, a MT stabilizing compound.

Using a fluorescent taxane analog combined with high resolution imaging of MT dynamics, it has recently been shown by Rai and coll. [26], that low non-saturating concentrations of a MT depolymerizing agent such as vinblastine, enhance catastrophes and induce a modulation of MT dynamics that favors an accumulation of PTX inside the MT. Such a mechanism could explain the observed synergy between Carba1 and PTX. To test if the same molecular mechanism is at work with Carba1, we first determined the Carba1 concentration able to induce catastrophes. We found that 10 μM Carba1 was able to induce a two-fold increase of the catastrophe frequency, which increased further to three-fold when combined with 100 nM PTX (**Fig 5G, Supplementary Fig 2A, B, C**). Further, Carba1 also increased the incorporation of a fluorescent derivative of PTX (Fchitax-3) within the MT shaft (**Fig 5H, I**). Such a result strongly suggests that the underlying mechanism for the observed synergy is that Carba1 binding to MTs induces a modification of the MT lattice leading to enhanced accumulation of PTX.

### Carba1 and PTX act synergistically to reduce tumor growth *in vivo*

Could the synergy between Carba1 and PTX, which we observed both at the molecular and the cellular levels, be translated into a therapeutic anti-cancer effect? To address this question, we compared the effects on tumor growth of Carba1 and PTX administrated separately to the effect of administration of Carba1 in combination with PTX, in a tumor mouse model. In a first series of experiments, we analyzed the effect of increasing doses of PTX or Carba1 when administered alone. To that aim, mice bearing sizable tumors, formed of HeLa cells that have been xenografted, received intravenous (i.v.) injections of PTX (from 2 to 8 mg/kg), every two days during 10 days (**Fig 6A**). In the same experiment, we analyzed the effect of Carba1 (from 15 to 60 mg/kg, i.v.) injected with the same schedule (**Fig 6B**). We found that PTX, when administered at 4 and 8 mg/kg, induced an important reduction of tumor size (**Fig 6A**). Carba1 did not induce a significant effect on tumor size whatever the dose injected, although a tendency towards smaller tumors appears with increasing Carba1 concentrations (**Fig 6B**). The results confirmed the anti-tumor effect of high PTX concentrations in this model. They also indicate that Carba1, when applied alone, has no significant anti-tumor activity, even at high concentrations. As shown in **supplementary Fig 3** the weight of PTX or Carba1 treated animals and vehicle-treated animals were not significantly different. Moreover the animals did not show any sign of discomfort, indicating a good tolerance of the treatments.

**Figure 6:**
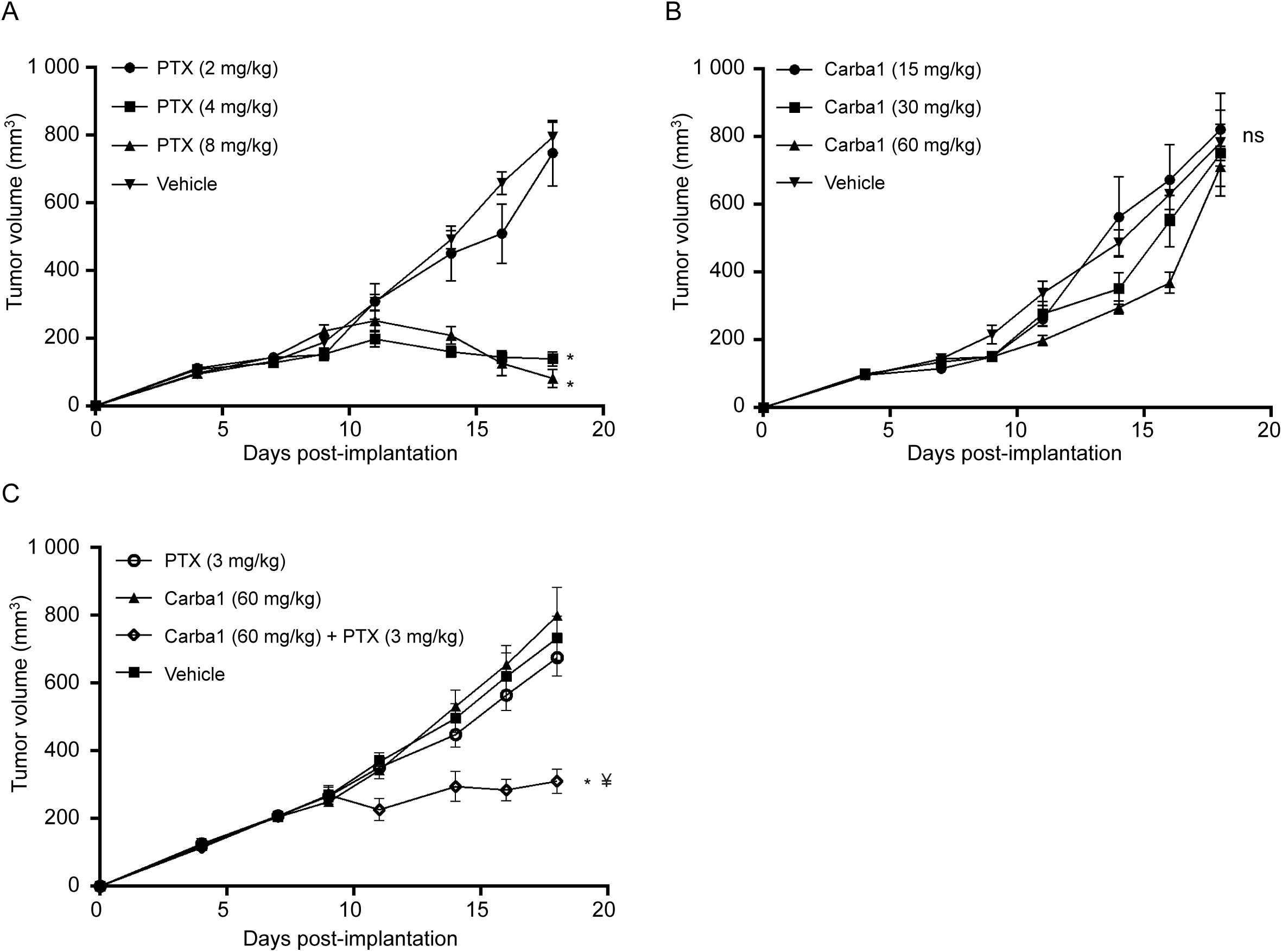
Effect of Carba1, PTX and their combination on tumor growth *in vivo*. A PTX inhibits the growth of HeLa cells xenografted in mice. When the tumors have reached a volume of about 200 mm^3^, mice were treated with PTX (2, 4 and 8 mg/kg) or the vehicle. Tumor growth was monitored with a sliding caliper. Error bars = SEM, * *p* < 0.01 compared to vehicle (ANOVA), n = 6 mice per group. B Carba1 has no significant effect on the growth of HeLa cells xenografted in mice. When the tumors have reached a volume of about 200 mm^3^, mice were treated with Carba1 (15, 30 and 60 mg/kg) or the vehicle. Tumor growth was monitored with a sliding caliper. Error bars = SEM, ns = non-significant, n = 6 mice per group. C The combination of otherwise ineffective doses of Carba1 and PTX inhibits the growth of HeLa cells xenografted in mice. When the tumors have reached a volume of about 200 mm^3^, mice were treated with PTX (3 mg/kg), Carba1 (60 mg/kg), the vehicle or the combination of PTX (3 mg/kg) and Carba1 (60 mg/kg). Tumor growth was monitored with a sliding caliper. Error bars = SEM, * *p* < 0.01 compared to PTX (3 mg/kg), ¥ *p* < 0.01 compared to Carba1 (60 mg/kg) (ANOVA), n = 8 mice per group.

We then conducted a study of the effect on tumor size of a low PTX dose (3 mg/kg) in combination with Carba1 (60 mg/kg). No modification of body weight was observed throughout the study, suggesting that the combination is well tolerated (**Supplementary Fig 4**). In this experiment PTX (3 mg/kg) has no effect on tumor size (**Fig 6C**). Compared to the previous experiment (no effect at 2 mg/kg, significant effect at 4 mg/kg), an intermediate effect would have been expected at 3 mg/kg. This indicates that the effect of PTX on tumors in this *in vivo* model should be observed at a threshold dose between 3 and 4 mg/kg.

As shown in **Fig 6C**, while no effect is observed when each compound is administered separately, a significant reduction in tumor size is observed with the combination of PTX and Carba1. These results indicate that the observed synergy between PTX and Carba1 *in vitro* also occurs *in vivo*, leading to an enhanced therapeutic efficacy at a low-dose of PTX treatments.

## Discussion

Our initial aim was to discover an agent that would allow lowering the dose of PTX while obtaining the same anti-tumor efficacy as the currently used therapeutic dose of PTX resulting in less toxicity. We thus screened a chemical library to detect compounds able to sensitize cells to a low, non-toxic dose of PTX. The test we used was a cytotoxicity test, therefore probing all vital cell functions. Whereas such a whole cell-based assay screens molecules having multiple potential targets and allows the biology to dictate the best targets [27], it may not be insignificant to have selected Carba1, an agent that targets tubulin and impairs MT dynamics. Indeed, this indicates that the most sensible target, in this specific context, is tubulin.

Recently, a series of carbazole-based MT targeting agents with anti-tumor properties has been reported, confirming the ability of this type of chemical scaffold to interact with the colchicine site of tubulin [28]. The Carba1 scaffold is a versatile one and we are currently synthesizing modified analogs for medicinal chemistry optimization.

The PTX binding site at the interior of the MT has been characterized at the atomic level: PTX binds to a pocket in β-tubulin that faces the MT lumen and is near the lateral interface between protofilaments (for review see [1]). The binding of PTX results in the expansion of the taxane binding pocket [29] of the tubulin dimer. Moreover PTX binding inhibits, in the protofilament, the compaction at the longitudinal interdimer interface, induced by GTP hydrolysis [30]. This allosteric mechanism would strengthen the longitudinal tubulin contacts leading to a stabilization of the MTs [1]. In this context, it is conceptually counterintuitive that an agent that depolymerizes MTs acts in synergy with PTX, an agent that stabilizes them. A possibility is that the binding of Carba1 to the tubulin dimer modifies its affinity for PTX. However, although it has been shown that the covalent occupancy of the taxane site can affect the structure of the colchicine site [31], the reverse has not yet been described. Moreover, in cells, due to the low affinity of Carba1 for tubulin and the nanomolar concentration of PTX that was used, it can be assumed that the probability that a single tubulin dimer has both a molecule of Carba1 and another of PTX bound is very low. Thus an allosteric effect at the level of the tubulin dimer, due to such a simultaneous binding, cannot be responsible for synergistic cytotoxicity.

Another possibility is that the binding of Carba1 can induce conformational changes of the growing MT ends that can facilitate the subsequent binding of PTX to the MT lattice. Recently, using TIRF analysis, it has been shown that inhibition of polymerization due to non-saturating doses of vinblastine induces a switch to catastrophes, which converts the MT plus end to a state that allows more accumulation of fluorescent PTX, indicating a higher affinity of MT growing ends for PTX [26]. Indeed, we have conducted the same type of experiment, replacing vinblastine with Carba1 and observed an increase in the rate of catastrophes associated with a greater incorporation of fluorescent PTX. Although the underlying structural mechanism is yet unknown, it is highly probable that Carba1 acts similarly to vinblastine to facilitate PTX accumulation.

It is known that PTX accumulates intracellularly [4], reaching intra-tumor concentrations that are higher in the tumors than in the plasma [7]. It is thus remarkable that the synergistic effect is observed not only at the MT level, but also at the cellular level, as well as when both drugs are applied systemically in animals to exert their anti-tumor action. Although it cannot be excluded that Carba1 has other targets, it is highly probable that the same mechanism is at work in these different contexts.

Because the combined administration of Carba1 and a low dose of PTX can have an anti-tumor effect equivalent to the administration of high doses of PTX alone, one could imagine that the combination should reduce the unwanted side effects of PTX. This has to be tested. For instance the effect of the combination should be compared to the PTX effect on the kinesin-based anterograde transport, since perturbation upon PTX treatment is thought to be part of the mechanism involved in peripheral neuropathy. But, given the mode of action we have described, with Carba1 facilitating the accumulation of PTX in MTs, we can bet that the combination should diminish MT-independent adverse events.

Anti-cancer strategies based on the concomitant administration of taxanes and depolymerizing agents such as vinorelbine have been reported [32–34]. However, these approaches used high doses of each of these drugs. Our results suggest that good therapeutic efficacy could be achieved with the combined administration of each of these agents at low doses, which could improve patient comfort. This work thus paves the way to new therapeutic perspectives that are easy to implement.

## Materials and methods

### Chemical Reagents and cells

Carba1 was synthesized at the CERMN (Centre d’Études et de Recherche sur le Médicament de Normandie, University of Caen). It was dissolved in anhydrous dimethyl sulfoxide (DMSO, Sigma, #D4540) and stored at −20°C as 10 mM stock solution. Paclitaxel (PTX) was purchased from Sigma (#T7402) and was dissolved in DMSO and stored at −20°C as 1 mM stock solution.

HeLa cells were obtained from the American Type Culture Collection (ATCC), routinely tested and authenticated by the ATCC. HeLa Kyoto cells expressing EGFP-alpha-tubulin and H2B-mcherry were from Cell Lines Service, #300670. Cells were grown in RPMI 1640 medium (Gibco, Invitrogen) supplemented with 1% penicillin/streptomycin and 10% Fetal Bovine Serum, and maintained in a humid incubator at 37°C in 5% CO_2_.

### Analysis of cell viability using MTT (screening of the chemical library)

The assay was performed in 96-well microplates. Cells were seeded at a density of 2,500 cells per well and allowed to adhere for 24 hours before being treated for 48 hours with either DMSO (0.1 % final concentration) or compounds at 5 μM, with or without 1 nM PTX. Viability was evaluated with a 3-(4,5-dimethylthiazol-2-yl)-2,5-diphenyl-tetrazolium bromide (MTT) colorimetric assay (Sigma, #M5655).

### Analysis of cell viability using Prestoblue assay

Cell viability was analyzed using the colorimetric Prestoblue assay (Invitrogen, #A13262). Cells were seeded in 96-well microplates (Greiner, #655077) at a density of 2,500 cells per well and allowed to adhere for 24 hours before being treated for 72 hours with either DMSO (0.1 % final concentration) or drugs at indicated concentrations. After a 72-hour treatment, 11 μL Prestoblue was added to each well and cells were incubated for another 45 minutes. The absorbance of each well was measured using a FLUOstar Optima microplate reader (Excitation, 544 nm; Emission, 580 nm).

### Apoptosis assay

The apoptosis assay was performed with FITC Annexin V Apoptosis Detection Kit I (BD Biosciences, #556547) using flow cytometry and analyzed by FCS express software.

### Cell cycle analysis

Cells were harvested and washed by centrifugation in PBS. Then, 10^5^ cells were fixed in 1 mL of 70% ethanol at 4°C overnight. Following two washes with PBS the cells were incubated with 50 μg/mL propidium iodide and 0.2 mg/mL RNase A (Sigma, #10109142001) / PBS for 30 minutes at 37°C before analysis. The percentage of cells in the specific cell-cycle phases (G0, G1, S, G2, and M) was determined using an Accuri C6 flow cytometer (Becton Dickinson).

### Immunofluorescence microscopy and live cell imaging

HeLa cells at a density of 20,000 cells were grown for 48 hours on glass coverslips placed in a 24-well microplate. When cells reached 70% confluence the medium was replaced with a fresh one supplemented with Carba1. After a 5-hour exposure to Carba1, cells were fixed and permeabilized with −20°C absolute methanol for 6 minutes. After washing and saturation with 3% BSA (Bovine Serum Albumin; Sigma, #A7906) / PBS (Phosphate Buffered Saline; Dutscher, #L0615-500), cells were incubated for 45 minutes at room temperature (RT) with anti-alpha-tubulin antibody (clone α3A1, 1:4000), produced by L. Lafanechère [13]. Cells were washed twice again and subsequently incubated with Alexa 488 conjugated anti-mouse antibody (1:1000, Jackson immunoresearch, #115-545-4637) for 30 minutes at RT. DNA was stained with 20 μM Hoechst 33342 (Sigma, #23491-52-3) and coverslips were mounted on glass slides with Mowiol 4-88 (Calbiochem, #475904). Images were captured with a Zeiss AxioimagerM2 microscope equipped with the acquisition software AxioVision and analyzed using the Fiji software. For live-cell imaging, HeLa Kyoto cells expressing EGFP-alpha-tubulin and H2B-mcherry were seeded on 2-well glass-slides (Ibidi, #80297) at a density of 7,000 cells per well and allowed to grow for 24 hours prior to imaging. After treatment, the slide was placed on a 37°C heated stage, at 5% CO_2_, and images were acquired every 2.5 minutes by a spinning disk confocal laser microscope (Andromeda iMIC) equipped with a Plan-Apochromat 20×/0.75 WD610 objective and an EMCCD camera (iXon 897). For each time point, a stack of 7 planes (thickness: 1 μm) was recorded. Acquisition (LA), off-line analysis (OA) and Fiji software programs were used.

### Transfection of GFP-EB3

To label MT plus ends, GFP-EB3 plasmids were used because EB3 has a strong binding affinity to MT plus ends. Cell transfection was performed using electroporation (AMAXA®, Köln, Germany). 2 μg of purified plasmid DNA were used for each transfection reaction.

### Fluorescence time-lapse videomicroscopy of MT plus ends

Live imaging of MT plus ends was performed as described in Honoré et al. [14], on transiently GFP-EB3 transfected-HeLa cells by using an inverted fluorescence microscope (ZEISS Axiovert 200M with a 63X objective). Time-lapse acquisition was performed with a COOLSNAP HQ (Roper Scientific), driven by Metamorph software (Universal Imaging Corp.). Image acquisition was performed at a temperature of 37 ± 1°C / 5% CO_2_ Data are from 3 independent experiments. For each experiment, 6 MTs/cell in 6 cells per condition were analyzed.

### Dynamic instability parameter analysis

The dynamic instability parameter analysis was performed by tracking MT plus ends over time using the imageJ software. The methods of calculation were as described in Honoré et al. [14].

### Tubulin Polymerization Assay

Tubulin was prepared from bovine brain as previously described [15]. Tubulin polymerization assays were carried out at 37°C in BRB80 buffer (80 mM Pipes, 0.5 mM MgCl_2_, 2 mM EGTA, 0.1 mM EDTA, pH 6.8 with KOH) by mixing 7 μM of pure tubulin, 1 mM GTP, 5 mM MgCl_2,_ and indicated concentrations of drugs (0.2% DMSO, final concentration) in a final volume of 100 μL. The time course of the self-assembly activity of tubulin was monitored as turbidity at 350 nm, 37°C, during 30 minutes, using a spectrophotometer (ThermoScientific, Evolution 201).

### [^3^H]-Colchicine Tubulin-Binding Assay

The tubulin was prepared from bovine brain as previously described [15]. Pure tubulin (3 μM final concentration) in cold BRB80 buffer was mixed at 4°C with [^3^H]-colchicine (82.6 Ci/mmol, Perkin-Elmer, #NET189250UC, 50 nM final concentration) and the competitor Carba1 (100 μM final concentration) in a final volume of 200 μL. Following a 30-minute incubation at 30°C, the samples were deposited onto 50 μL of presedimented DEAE Sephadex A25 in BRB80 buffer. All subsequent steps were carried out at 4°C. Samples were incubated for 10 minutes with continuous shaking to ensure quantitative binding of tubulin to the gel. Following centrifugation (2400*g*, 4 minutes), supernatants were discarded and the pellets containing the bound molecule-tubulin complexes were washed four times with 1 mL of BRB80 buffer. Pellets were incubated for 10 minutes with 500 μL of ethanol to solubilize the tubulin-bound tritiated colchicine and 400 μL aliquots of the ethanol solutions were transferred to 5 mL of Ultima Gold scintillant (Perkin-Elmer) for determination of radioactivity.

### Determination of the binding constant of Carba1 on tubulin using a competition assay

Calf brain tubulin was purified as described [16]. 2-Methoxy-5-(2,3,4-trimethoxyphenyl)-2,4,6-cycloheptatrien-1-one (MTC)[17] was a kind gift of Prof. T. J. Fitzgerald (School of Pharmacy, Florida A & M University, Tallahassee, FL). The compounds were diluted in 99.8% DMSO-d6 (Merck, Darmstadt, Germany) to a final concentration of 10 mM and stored at −80°C.

Competition of the compound with MTC was tested by the change in the intensity of fluorescence of MTC upon binding to tubulin. The fluorescence emission spectra (excitation at 350 nm) of 10 μM tubulin and 10 μM MTC in 10 mM sodium phosphate, 0.1 mM GTP, pH 7.0, were measured in the presence of different concentrations (0 to 20 μM) of the desired ligand with 5 nm excitation and emission slits using a Jobin-Ybon SPEX Fluoromax-2 (HORIBA, Ltd., Kyoto, Japan). The decrease in the intensity of the fluorescence in the presence of the competitor ligand indicated competition for the same binding site. The data were analyzed and the binding constants determined using Equigra V5.0 as described in Díaz and Buey [18].

### In vitro MT dynamics and analysis of MT dynamics parameters

Perfusion chambers were obtained by assembling silane-PEG-biotin (LaysanBio, #MW3400) coverslips and silane-PEG (Creative PEGWork, #PSB-2014) glass slides as described previously [15]. The chambers were perfused with Neutravidin (25 μg/mL in 1% BSA, ThermoFisher Scientific, #31000), PLL-PEG (0.1 mg/mL in 10 mM HEPES, pH 7.4, Jenkem, #PLL20K-G35-PEG2K), 1% BSA in BRB80, and GMPCPP-stabilized (Jena Bioscience, #NU-405S), ATTO-488-labeled MT seeds (ATTO-Tec). MT assembly was initiated with 12 μM tubulin (containing 20% ATTO 488-labeled tubulin) in the presence of 4 μM Carba1. Time-lapse imaging was performed on an inverted Eclipse Ti (Nikon) microscope with an Apochromat 60×/1.49 numerical aperture (NA) oil immersion objective (Nikon) equipped with an ilas2 TIRF system (Roper Scientific). We performed time-lapse imaging at 1 frame per 2 seconds with an 80 milliseconds exposure time. MT dynamics parameters were determined on kymographs using ImageJ software. *In vitro* assay for MT growth dynamics and analysis of MT dynamic parameters in the presence of tubulin (cytoskeleton) and EB3 with Carba1 and Fchitax-3 was performed as described previously [26]. For statistical analysis, graphs were plotted in GraphPad Prism 7 and statistical analysis was done using non-parametric Mann-Whitney U-test.

### Tumor xenografts in mice

All animal studies were performed in accordance with the institutional guidelines of the European Community (EU Directive 2010/63/EU) for the use of experimental animals and were authorized by the French Ministry of Higher Education and Research under the reference: apafis#8854-2017031314338357 v1.

In a first series of experiment, the effects of PTX or Carba1 when administrated alone were evaluated. To that aim anesthetized (4% isoflurane/air for anesthesia induction and 1.5% thereafter) five-week-old female NMRI nude mice (Janvier Labs, Le Genest-Saint Isle, France) were injected subcutaneously in the flank with 10^7^ exponentially dividing Hela cells in 1X PBS. Tumor size was measured three times a week using a caliper, and the tumor volume was calculated as follows: length × (width)^2^ × 0.4. When tumors have reached a volume of about 200 mm^3^ i.e. nine days after cell injection, mice were randomized in 7 groups of 6 mice each and drugs were injected intravenously every two days. A first group received the vehicle (14% DMSO, 14% Tween 80 and 72% PBS). Three groups received PTX at different doses (2, 4 and 8 mg/kg) while three other groups received Carba1 at different doses (15, 30 and 60 mg/kg).

In a second series of experiments, the effect of a combination of PTX-Carba1 was evaluated, and compared to the effect of the compounds alone. To that aim, five-week-old female NMRI nude mice were injected subcutaneously with 10^7^ exponentially dividing HeLa cells into the right flank. When tumors have reached a volume of about 200 mm^3^ i.e. nine days after cell injection, mice were randomized in 4 groups of 8 mice each and drugs were injected intravenously every two days. The first group received PTX at 3 mg/kg, the second group Carba1 at 60 mg/kg, the third group received a combination of Carba1 (60 mg/kg) and PTX (3 mg/kg), and the fourth group received the vehicle (14% DMSO, 14% Tween 80 and 72% PBS). Groups were statistically compared using ANOVA.

## Acknowledgments

This work was supported by INSERM, Université Grenoble Alpes, CNRS, and by l’Institut National du Cancer (INCa, PLBIO16186), Fondation ARC (PJA 20151203348) and Association “Le Cancer du Sein, Parlons-en!”, to LL. We thank Isabelle Arnal and Christian Delphin for their help in the purification of tubulin and in the TIRF experiments, Ganadería Fernando Díaz for calf brain supply, Wei-Shuo Fang for synthesizing Fchtitax-3 and Anne Martinez for her help in the initial step of the study. We also thank the Developmental therapeutics program at the National Cancer Institute for the screening of Carba1 on 60 cancer cell lines.

## Author contributions

L.L. conceived the project and supervised the findings of this work. L.P. designed and realized in cellulo and in vitro experiments under the supervision of L.L., E.D. and A. And. A. And. and E.D. advised on the mechanism of action. R.P. conceived and designed the screening and A.V. performed the screening and its analysis. A.R. performed Fichtax-3 experiments and analysis and contributed to Fig. 5. S.R.R. realized tubulin purification and contributed to the experiments measuring microtubule dynamics. S.M. and A.S.R. performed additional cell cytotoxicity experiments. D.L.A., M.A.O. and J.F.D. performed the analyses of the affinity of Carba1 for tubulin and contributed to Fig. 6. M.G. and J.V. realized the animal experiments, their statistical analysis and contributed to Fig. 6.

V.J. and J.L.C conceived the animal experiments and interpreted the data.

P.S. and P.D. synthesized and provided the chemical compounds.

A. Akh., J.F.D, E.D., A. And. and K.S. contributed to the interpretation of the results.

L.L., L.P. and K.S. wrote the manuscript and designed the figures with input of all the authors.

## The Paper Explained

### PROBLEM

Paclitaxel (Taxol®) is a drug that has been proven in cancer chemotherapy. However, its administration poses problems of toxicity, undesirable side effects and resistance.

To overcome this problem, rather than looking for new drugs with the same mechanism of action as paclitaxel, but less toxic, we looked for drugs that work synergistically with paclitaxel to kill cancer cells, by screening a large chemical library. The underlying idea was to find a way to achieve the same therapeutic efficacy, but with lower doses of paclitaxel.

### RESULTS

We describe a compound that acts synergistically with paclitaxel. This compound acts using a recently described mechanism: it modulates the dynamics of the end of the microtubules, facilitating the accumulation of paclitaxel inside the microtubule. This action at the microtubule level results in reduced tumor growth in an animal model. Thus, the same effectiveness as a therapeutic dose of paclitaxel is obtained when a lower dose of paclitaxel is used in combination with the compound.

### IMPACT

The presented results pave the way for new therapeutic strategies, based on the combination of low doses of microtubule targeting agents with opposite mechanisms of action. Such combinations may reduce toxicity and adverse side effects due to high doses of microtubule targeting agents used in current treatments.

## References

1. Steinmetz MO, Prota AE (2018) Microtubule-Targeting Agents: Strategies To Hijack the Cytoskeleton. Trends Cell Biol 28:776–792.

2. Akhmanova A, Steinmetz MO (2015) Control of microtubule organization and dynamics: two ends in the limelight. Nat Rev Mol Cell Biol 16: 711–726.

3. Dumontet C, Jordan MA (2010) Microtubule-binding agents: a dynamic field of cancer therapeutics. Nat Rev Drug Discov 9: 790–803.

4. Jordan MA, Toso RJ, Thrower D, Wilson L (1993) Mechanism of mitotic block and inhibition of cell proliferation by taxol at low concentrations. Proc Natl Acad Sci U S A 90: 9552–9556.

5. Walsh V, Goodman J (2002) From taxol to taxol®: The changing identities and ownership of an anti-cancer drug. Med Anthropol 21: 307–336.

6. Mitchison TJ (2012) The proliferation rate paradox in antimitotic chemotherapy. Mol Biol Cell 23: 1–6.

7. Zasadil LM, Andersen KA, Yeum D, Rocque GB, Wilke LG, Tevaarwerk AJ, Raines RT, Burkard ME, Weaver BA (2014) Cytotoxicity of paclitaxel in breast cancer is due to chromosome missegregation on multipolar spindles. Sci Transl Med 6: 229ra43.

8. Komlodi-Pasztor E, Sackett D, Wilkerson J, Fojo T (2011) Mitosis is not a key target of microtubule agents in patient tumors. Nat Rev Clin Oncol 8: 244–250.

9. Mitchison TJ, Pineda J, Shi J, Florian S (2017) Is inflammatory micronucleation the key to a successful anti-mitotic cancer drug? Open Biol 7: 170182.

10. Kavallaris M (2010) Microtubules and resistance to tubulin-binding agents. Nat Rev Cancer 10: 194–204.

11. Millecamps S, Julien J-P (2013) Axonal transport deficits and neurodegenerative diseases. Nat Rev Neurosci 14: 161–176.

12. Smith JA, Slusher BS, Wozniak KM, Farah MH, Smiyun G, Wilson L, Feinstein S, Jordan MA (2016) Structural Basis for Induction of Peripheral Neuropathy by Microtubule-Targeting Cancer Drugs. Cancer Res 76: 5115–5123.

13. Peris L, Thery M, Faure J, Saoudi Y, Lafanechere L, Chilton JK, Gordon-Weeks P, Galjart N, Bornens M, Wordeman L, et al. (2006) Tubulin tyrosination is a major factor affecting the recruitment of CAP-Gly proteins at microtubule plus ends. J Cell Biol 174: 839–849.

14. Honore S, Braguer D (2011) Investigating microtubule dynamic instability using microtubule-targeting agents. Methods Mol Biol 777: 245–260.

15. Ramirez-Rios S, Serre L, Stoppin-Mellet V, Prezel E, Vinit A, Courriol E, Fourest-Lieuvin A, Delaroche J, Denarier E, Arnal I (2017) A TIRF microscopy assay to decode how tau regulates EB’s tracking at microtubule ends. Methods Cell Biol 141: 179–197.

16. Andreu JM (2007) Large scale purification of brain tubulin with the modified Weisenberg procedure. Methods Mol Med 137: 17–28.

17. Fltzgerald TJ (1976) Molecular features of colchicine associated with antimitotic activity and inhibition of tubulin polymerization. Biochem Pharmacol 25: 1383–1387.

18. DÍaz JF, Buey RM (2007) Characterizing Ligand-Microtubule Binding by Competition Methods. In, Methods in molecular medicine pp 245–260.

19. Mohan R, Katrukha EA, Doodhi H, Smal I, Meijering E, Kapitein LC, Steinmetz MO, Akhmanova A (2013) End-binding proteins sensitize microtubules to the action of microtubule-targeting agents. Proc Natl Acad Sci U S A 110: 8900–8905.

20. Shoemaker RH (2006) The NCI60 human tumour cell line anticancer drug screen. Nat Rev Cancer 6: 813–823.

21. Maiato H, Gomes A, Sousa F, Barisic M (2017) Mechanisms of Chromosome Congression during Mitosis. Biology (Basel) 6: 13.

22. Prudent R, Moucadel V, Nguyen C-HH, Barette C, Schmidt F, Florent J-CC, Lafanechère L, Sautel CF, Duchemin-Pelletier E, Spreux E, et al. (2010) Antitumor Activity of Pyridocarbazole and Benzopyridoindole Derivatives that Inhibit Protein Kinase CK2. Cancer Res 70: 9865–9874.

23. Prudent R, Vassal-Stermann E, Nguyen C-H, Pillet C, Martinez A, Prunier C, Barette C, Soleilhac E, Filhol O, Beghin A, et al. (2012) Pharmacological inhibition of LIM kinase stabilizes microtubules and inhibits neoplastic growth. Cancer Res 72: 4429–4439.

24. Jordan MA, Thrower D, Wilson L (1992) Effects of vinblastine, podophyllotoxin and nocodazole on mitotic spindles. Implications for the role of microtubule dynamics in mitosis. J Cell Sci 102: 401–416.

25. La Regina G, Edler MC, Brancale A, Kandil S, Coluccia A, Piscitelli F, Hamel E, De Martino G, Matesanz R, Díaz JF, et al. (2007) Arylthioindole Inhibitors of Tubulin Polymerization. 3. Biological Evaluation, Structure-Activity Relationships and Molecular Modeling Studies. J Med Chem 50: 2865–2874.

26. Rai A, Liu T, Glauser S, Katrukha EA, Estévez-Gallego J, Rodríguez-García R, Fang W-S, Díaz JF, Steinmetz MO, Altmann K-H, et al. (2019) Taxanes convert regions of perturbed microtubule growth into rescue sites. Nat Mater.

27. Peterson JR, Mitchison TJ (2002) Small molecules, big impact: a history of chemical inhibitors and the cytoskeleton. Chem Biol 9: 1275–1285.

28. Diaz P, Horne E, Xu C, Hamel E, Wagenbach M, Petrov RR, Uhlenbruck B, Haas B, Hothi P, Wordeman L, et al. (2018) Modified carbazoles destabilize microtubules and kill glioblastoma multiform cells. Eur J Med Chem 159: 74–89.

29. Kellogg EH, Hejab NMA, Howes S, Northcote P, Miller JH, Díaz JF, Downing KH, Nogales E (2017) Insights into the Distinct Mechanisms of Action of Taxane and Non-Taxane Microtubule Stabilizers from Cryo-EM Structures. J Mol Biol 429: 633–646.

30. Alushin GM, Lander GC, Kellogg EH, Zhang R, Baker D, Nogales E (2014) High-resolution microtubule structures reveal the structural transitions in αβ-tubulin upon GTP hydrolysis. Cell 157: 1117–1129.

31. Field JJ, Pera B, Gallego JE, Calvo E, Rodríguez-Salarichs J, Sáez-Calvo G, Zuwerra D, Jordi M, Andreu JM, Prota AE, et al. (2018) Zampanolide Binding to Tubulin Indicates Cross-Talk of Taxane Site with Colchicine and Nucleotide Sites. J Nat Prod 81:494–505.

32. Limentani SA, Brufsky AM, Erban JK, Jahanzeb M, Lewis D (2006) Phase II Study of Neoadjuvant Docetaxel/Vinorelbine Followed by Surgery and Adjuvant Doxorubicin/Cyclophosphamide in Women with Stage II/III Breast Cancer. Clin Breast Cancer 6: 511–517.

33. Tortoriello A, Facchini G, Caponigro F, Santangelo M, Benassai G, Persico G, Citarella A, Carola M, Marzano N, Iaffaioli RV (1998) Phase I/II study of paclitaxel and vinorelbine in metastatic breast cancer. Breast Cancer Res Treat 47: 91–97.

34. Berruti A, Bitossi R, Gorzegno G, Bottini A, Generali D, Milani M, Katsaros D, Rigault de la Longrais IA, Bellino R, Donadio M, et al. (2005) Paclitaxel, vinorelbine and 5-fluorouracil in breast cancer patients pretreated with adjuvant anthracyclines. Br J Cancer 92: 634–638.

